# Genomic sequencing of *Phyllosticta citriasiana* provide insight into its conservation and diversification with closely related *Phyllosticta* species associated with citrus

**DOI:** 10.1101/740852

**Authors:** Mingshuang Wang, Bei Liu, Ruoxin Ruan, Yibing Zeng, Jinshui Luo, Hongye Li

**Author notes:** Authors for Correspondence: (M. Wang), (H. Li).

## Abstract

*Phyllosticta citriasiana* is the causal agent of the pomelo tan spot. Here, we presented the ~34Mb genome of *P. citriasiana.* The genome is organized in 92 contigs, encompassing 9202 predicted genes. Comparative genomic analyses with other two *Phyllosticta* species (*P. citricarpa* and *P. capitalensis*) associated with citrus was conducted to understand their evolutionary conservation and diversification. Pairwise genome alignments revealed that these species are highly syntenic. All species encode similar numbers of CAZymes and secreted proteins. However, the molecular functions of the secretome showed that each species contains some enzymes with distinct activities. Three *Phyllosticta* species shared a core set of 7261 protein families. *P. capitalensis* had the largest set of orphan genes (2040), in complete contrast to that of *P. citriasiana* (371) and *P. citricarpa* (262). Most of the orphan genes were functionally unknown, but they contain a certain number of species-specific secreted proteins. A total of 23 secondary metabolites (SM) biosynthesis clusters were identified in the three *Phyllosticta* species, 21 of them are highly conserved among these species while the remaining 2 showed whole cluster gain and loss polymorphisms or gene content polymorphisms. Taken together, our study reveals insights into the genetic mechanisms of host adaptation of *Phyllosticta* species associated with citrus and paves the way to identify effectors that function in infection of citrus plants.

## Introduction

Citrus Black Spot (CBS), caused by *Phyllosticta citricarpa*, is an important disease of citrus, it can affect almost all grown citrus cultivars, including sweet orange *(Citrus sinensis)*, mandarin (*C. reticulata* and *C. unshiu)*, pomelo (*C. maxima)*, grapefruit (*C. paradise*) and lemon *(C.limon*) (Wang et al., 2012). This disease mainly causes black lesions in the fruits, making the fruits unsuitable for the fresh market. When the disease is severe, yield losses are significant due to premature fruit drop (Kotzé, 1981). CBS mainly happened in humid subtropical regions, including Asia, Africa, South America, Australia, and most recently, Florida (Kotzé, 1981; Wang et al., 2012; Miles et al., 2013; Wang et al., 2016b; Carstens et al., 2017). As this disease is previously absent in Mediterranean countries like Spain, Italy, Israel, and Turkey, *P. citricarpa* was listed as an A1 quarantine pest by European Union (Paul et al., 2005; EFSA, 2014). However, a recent survey reported the presence of *P. citricarpa* in Italy, Malta and Portugal (Guarnaccia et al., 2017).

Besides *P. citricarpa*, other species of *Phyllosticta* have been reported to be associated with citrus. *P. citriasiana*, first identified to be a harmful pathogen of pomelos in 2009, was able to cause necrotic spots (tan spots) on fruit similar to those caused by *P. citricarpa* (Wulandari et al., 2009). By performing multi-locus phylogenetic analyses on a large number of *Phyllosticta* species collected in China, Wang *et* al.(2012) found that *P. citriasiana* was isolated only from pomelos, and *P. citricarpa* was isolated from lemons, mandarins, and oranges, but never from pomelos, indicating that the citrus-associated pathogenic *Phyllosticta* population may have a host-related differentiation (Wang et al., 2012). In addition to *P. citriasiana*, many other *Phyllosticta* fungi were also found in citrus, such as *P. capitalensis, P. citribraziliensis, P.citrichinaensis, P. paracapitalensis*, and *P. paracitricarpa* (Glienke et al., 2011; Wang et al., 2012; Guarnaccia et al., 2017). Of them, *P. capitalensis* is the most frequently isolated species. This species has a very wide distribution and it has been isolated as endophytes from dozens of plants (Wikee et al., 2013).

Due to the early discovery of *P. citricarpa* causing CBS and its economic importance, *P. citricarpa* is extensively studied and many information is now available on this pathogen’s population structure, reproduction mode and introduction pathways (Spósito et al., 2011; Wang et al., 2016b; Carstens et al., 2017). However, little is known about the newly identified pathogen of pomelo tan spot, *P. citriasiana.* In this study, we sequenced the genome of *P. citriasiana*, generating a high-quality reference genome assembly and provide an overview of the genome structure of this important pathogen; we also compared its genome with other two closely related *Phyllosticta* species associated with citrus, i.e., *P. citricarpa* and *P. capitalensis* to provide insight into their evolutionary conservation and diversification.

## Materials and Methods

### Fungal strain

The *Phyllosticta citriasiana* strain ZJUCC200914 (CGMCC3.14344) was isolated from tan spot infected pomelos collected from Fujian Province, China (Wang et al., 2012). Cultures of strains were maintained on regular solid PDA (potato dextrose agar) or in liquid potato dextrose broth (PDB) at 25°C.

### Genome assembly and annotation

The genomic DNA and RNA of *P. citriasiana* were extracted from mycelia grown in PDB as described previously (Wang et al., 2015). The genome was first surveyed through Illumina HiSeq 2500 platform using TruSeq libraries (150bp paired-end reads, insert size of 350bp) and then sequenced using the long reads PacBio technology. A total of 1.9 Gb PacBio data, 6.7 Gb pair-end data were generated in the sequencing process, which corresponds to ~250 fold of sequence depth. To obtain high-quality gene calls, RNA-Seq was conducted with the same sample and 6.0Gb Illumina paired-end reads were obtained.

To obtain the *P. citriasiana* genome, the PacBio reads were initially assembled using Canu 1.7 (Koren et al., 2017) and error correction was conducted using Pilon version 1.22 with the Illumina reads (Walker et al., 2014). Genome quality assessment was performed through the presence of conserved single-copy fungal genes using BUSCO version 3 (Simao et al., 2015). RNA-seq data were aligned to the genome using Bowtie 2.3.4 and TopHat 2.0.9 (Langmead and Salzberg, 2012; Kim et al., 2013). Genome annotation was performed using the BRAKER version 1.0 pipeline combining the RNA-seq-based gene prediction and ab initio gene prediction (Hoff et al., 2015).

The genomes of *P. citricarpa* (accession number LOEO00000000.1) and *P. capitalensis* (accession number LOEN00000000.1) were downloaded from the NCBI genome database (Wang et al., 2016b). Gene model predictions of these two genomes were generated with AUGUSTUS 3.1 using the training annotation file of *Phyllosticta citriasiana* (Stanke et al., 2008). Repetitive elements were annotated in all assemblies using RepeatMasker version open-4.0.7 (http://www.repeatmasker.org). For pairwise syntenic analysis of genome structures, the contigs of the paired genomes of *Phyllosticta* species were aligned with the MUMmer 3.23 package (Delcher et al., 2003). The average nucleotide identity was estimated using the ANI calculator (Rodriguez-R and Konstantinidis, 2016). The statistical reports for genomes were calculated by using in-house Perl scripts.

### Functional annotation of genes

To functionally annotate gene models, we assigned protein sequence motifs to protein families (Pfam) and gene ontology (GO) terms using the Pfam and eggNOG databases (Huerta-Cepas et al., 2017; El-Gebali et al., 2018). The GO enrichment in molecular functions was produced with the dcGO database (Fang and Gough, 2013). Protein orthogroups were clustered using the orthoMCL algorithm in combination with an all-versus-all protein BLAST search (e-value < 1e-10, identity > 50%) (Chen et al., 2006). The carbohydrate-active enzymes were annotated by the web-based dbCAN2 meta server (Zhang et al., 2018). To identify secreted proteins, we use SingalP 4.1 to predict the transmembrane domains and we excluded non-extracellular and GPI-anchored proteins by using targetP 1.1 (Emanuelsson et al., 2000) and GPI-SOM (Fankhauser and Mäser, 2005). Fungal secondary metabolite pathways were predicted using the online tool antiSMASH 4.0 (Blin et al., 2017).

### Data availability

The assembled *Phyllosticta citriasiana* genome has been deposited in GenBank under the accession number QOCM00000000. All the annotation data generated in this study have been deposited on the figshare repository at DOI: 10.6084/m9.figshare.9178061 (the data will be made publicly available upon acceptance of the manuscript).

## Results and discussion

### Genome assembly and general features

We assembled the genome of *P. citriasiana* using a combination of Illumina and PacBio reads with ~250 fold of sequence depth. The de novo assembly resulted in a genome size of 34.2 Mb assembled in 92 contigs with an N50 of ~1Mb. The genomes of *P. citricarpa* and *P. capitalensis* previously sequenced were utilized and annotated in this study (Wang et al., 2016b). The completeness of these three genome assemblies was estimated by BUSCO (Simao et al., 2015). We found 1759 out of 1732 (98.4%) BUSCO groups were identified in the *P. citriasiana* genome, indicating a high degree of completeness. Although the assembly of *P. citricarpa* and *P. capitalensis* possess a large number of contigs, the BUSCO results showed that they are around 95% completeness, suggesting that these genomes are reliable for the downstream analyses (Table 1).

**Table 1.** Genomic features of three *Phyllosticta* species associated with citrus.

| Features | *P. capitalensis* strain | *P. citricarpa* strain | *P. citriasiana* strain |
| --- | --- | --- | --- |
|  | Gm33 | Gc12 | CGMCC3.14344 |
| Genome size (Mb) | 32.4 | 31.1 | 34.2 |
| BUSCOs (%) | 95.5 | 94.8 | 98.4 |
| Number of contigs | 1341 | 5748 | 92 |
| N50 (Kb) | 20.8 | 76 | 968.9 |
| GC content (%) | 54.6 | 53.1 | 51.4 |
| Protein-coding genes | 9983 | 9083 | 9202 |
| Gene density (number of genes per Mb) | 308 | 292 | 269 |
| Mean gene length (bp) | 1632 | 1642 | 1677 |
| Repeat rate (%) | 0.29 | 0.97 | 2.19 |

The overall G + C content of *P. citriasiana* (51.4%) is apparently lower than that of *P. citricarpa* (53.1%) and *P. capitalensis* (54.6%). The percentage of repetitive sequences of *P. citriasiana* is 2.19%, around 2-fold of that of *P. citricarpa* (0.97%) and 7-fold of *P. capitalensis* (0.29%). *P. citriasiana* has the lowest gene density but the longest gene length among the three species. The genome of *P. citriasiana* was predicted to have 9202 proteins, which is comparable with that of *P. citricarpa* (9083), but much lower than that (9983) of *P. capitalensis* genome (Table 1).

During preparing this manuscript, we noticed that a paper describing the genomic sequencing of *P. citricarpa* and *P. capitalensis* was published (Rodrigues et al., 2019). However, the general features of their genome sequences are of great difference from ours. For example, the authors predicted ~15000 proteins for both species while our data only predicted ~9500 ones. In that study, we found that the *P. citricarpa* genomic assembly consisted of 19,143 contigs with the N50 of 3049bp and the *P. capitalensis* genomic assembly contains 11,080 contigs with the N50 of 4925bp (Rodrigues et al., 2019). This means that their genome sequence was very fragmented and the incompleteness of the genome was also confirmed by their BUSCO analysis (Rodrigues et al., 2019). Thus, comparing with their genomic data, we believe that the data in this study is much better and more reliable.

### Genomic similarity

The pairwise comparison analysis based on oriented contigs reveals a high degree of genome-wide macrosynteny between *P. citriasiana* and the other two species (Fig. 1A). However, the average nucleotide identity (95.98%) between *P. citriasiana* and *P. citricarpa* is much larger than that (81.19%) between *P. citriasiana* and *P. capitalensis* (Fig. 1B), indicating that *P. citriasiana* is much closer to *P. citricarpa* and relatively far from *P. capitalensis*.

**Fig. 1.**
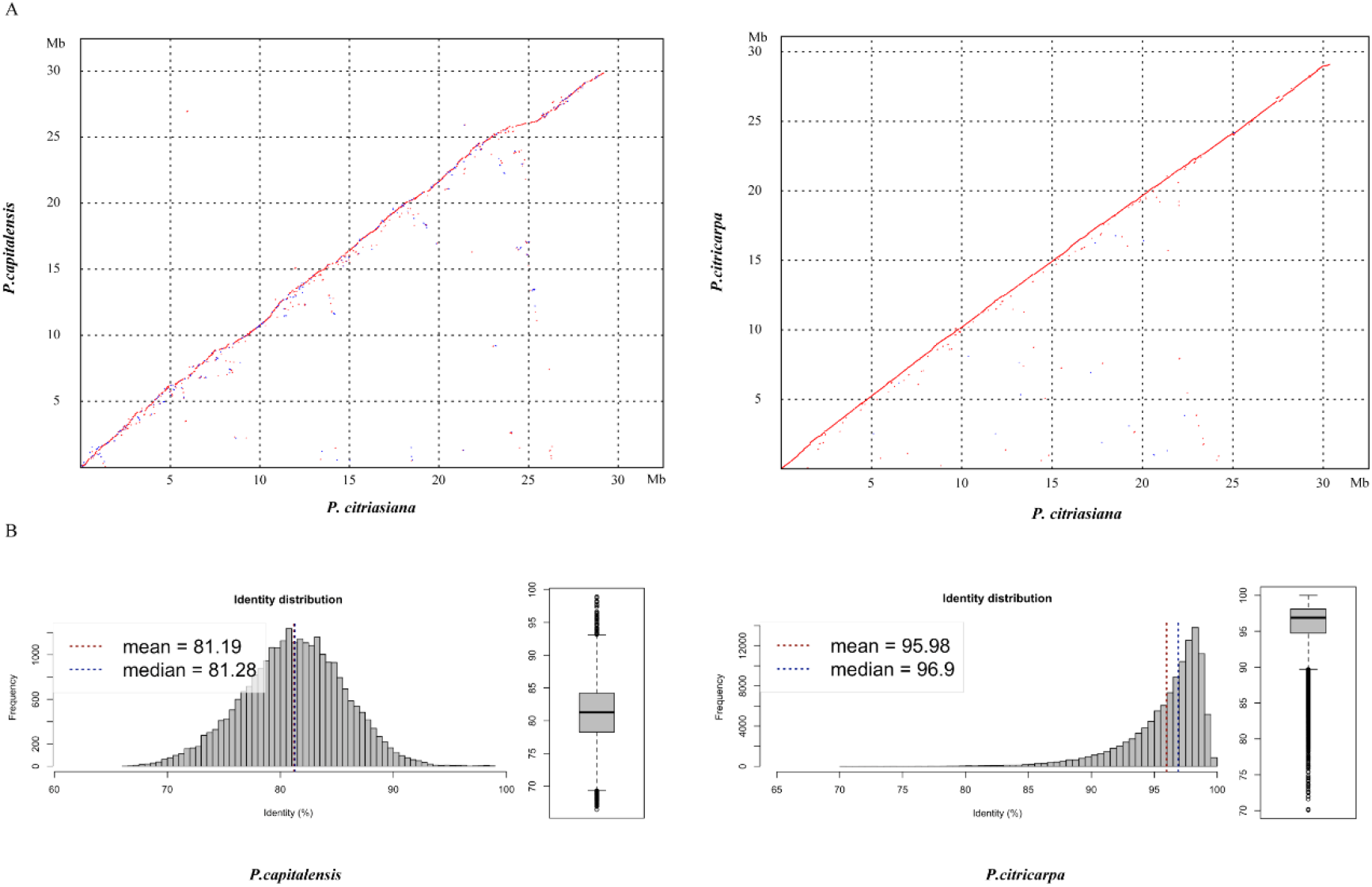
Genomic similarity between *P. citriasiana* and the other two *Phyllosticta* species (*P. citricarpa* and *P. capitalensis*) associated with citrus. **A)** Dotplos showing genome nucleotide alignments of *P. citriasiana* with *P. citricarpa* and *P. capitalensis.* Red diagonals represent alignments in the same direction, whereas blue ones suggest a reverse orientation. **B)** Distribution of nucleotide identities between *P. citriasiana* and the other two *Phyllosticta* species.

### Carbohydrate active enzymes and secretomes

The cell wall in plant forms a complex network of different polysaccharides that includes cellulose, hemicellulose, pectin, and lignin. Carbohydrate-active enzymes (CAZymes) play important roles in the breakdown of complex carbohydrates and are responsible for the acquisition of nutrients from the plant for plant-associated fungi (Kubicek et al., 2014). A total of 183 putative CAZyme genes were identified in *P. citriasiana*, which includes 100 Glycoside Hydrolases (GHs), 46 Glycosyl Transferases (GTs), 30 Auxiliary Activities (AAs), 3 Carbohydrate Esterases (CEs), 3 Polysaccharide Lyases (PLs), and 1 Carbohydrate-Binding Modules (CBMs) (Table S1-4). The types and numbers of CAZymes among different species of *Phyllosticta* are very similar (Table S1-4). When compared with other species in the Dothideomycetes, *P. citriasiana* appears to contain a much less extensive set of CAZymes, for example, *Alternaria alternata* has 373 CAZymes while *Zymoseptoria tritici* contains 324 ones (Goodwin et al., 2011; Ohm et al., 2012; Wang et al., 2016a). The smaller number of CAZymes in *Phyllosticta* coincides with the phenomenon that these species have a relatively long time to infect citrus fruits and the scab expanded slowly in the fruit peels (Wang et al., 2012; Goulin et al., 2016).

Pathogens can secrete a series of proteins that are deployed to the host-pathogen interface during infection, and secretome proteins play an important role in pathogenicity (Presti et al., 2015). Approximately 5% of the total proteins (470) of *P. citriasiana* are predicted to be secreted (Table S5-8). ‘Hydrolase activity’ was the most abundant molecular function of the secretome, other GO terms over-represented among the secreted proteins include ‘hydrolase activity, acting on glycosyl bonds’, ‘carboxylic ester hydrolase activity’, ‘lipase activity’, ‘exopeptidase activity’, ‘serine hydrolase activity’, and ‘hydrolase activity, acting on carbon-nitrogen bonds’ (Table 2). The other two *Phyllosticta* species contain a similar number of SPs with *P. citriasiana*, i.e., 465 in *P. citricarpa* and 491 in *P. capitalensis* (Table S5-8). However, their GO categories showed some differences from that of P. *citriasiana.* The SPs of *P. citricarpa* were not enriched in ‘exopeptidase activity’, ‘serine hydrolase activity’, and ‘hydrolase activity, acting on carbon-nitrogen bonds’ but was in ‘transferase activity, transferring hexosyl groups’ (Table 2). The SPs of *P. capitalensis* lacks ‘exopeptidase activity’ and ‘hydrolase activity, acting on carbon-nitrogen bonds’ but contains ‘phosphatase activity’, ‘transferase activity, transferring hexosyl groups’, and ‘UDP-glycosyltransferase activity’ (Table 2). These results suggested that the constitution of different *Phyllosticta* secretomes has changed, each species have some preferred enzymes with distinct activities.

**Table 2.** Enriched molecular functional categories for secreted proteins genes of *Phyllosticta* species associated with citrus.

| Species | GO id | GO term | FDR | Gene number |
| --- | --- | --- | --- | --- |
| *P. citriasiana* | GO:0016787 | hydrolase activity | 2.82E-21 | 82 |
|  | GO:0016798 | hydrolase activity, acting on glycosyl bonds | 1.34E-19 | 29 |
|  | GO:0052689 | carboxylic ester hydrolase activity | 4.44E-07 | 12 |
|  | GO:0016298 | lipase activity | 3.61E-05 | 10 |
|  | GO:0008238 | exopeptidase activity | 8.67E-04 | 8 |
|  | GO:0017171 | serine hydrolase activity | 6.66E-03 | 7 |
|  | GO:0016810 | hydrolase activity, acting on carbon-nitrogen bonds | 9.33E-03 | 8 |
| *P. citricarpa* | GO:0016787 | hydrolase activity | 4.80E-17 | 76 |
|  | GO:0016798 | hydrolase activity, acting on glycosyl bonds | 1.57E-20 | 30 |
|  | GO:0052689 | carboxylic ester hydrolase activity | 4.56E-06 | 11 |
|  | GO:0016298 | lipase activity | 2.10E-04 | 9 |
|  | GO:0016758 | transferase activity, transferring hexosyl groups | 2.42E-03 | 10 |
| *P. capitalensis* | GO:0016787 | hydrolase activity | 1.48E-18 | 100 |
|  | GO:0016798 | hydrolase activity, acting on glycosyl bonds | 6.07E-15 | 29 |
|  | GO:0052689 | carboxylic ester hydrolase activity | 5.68E-08 | 15 |
|  | GO:0016298 | lipase activity | 1.76E-04 | 11 |
|  | GO:0016791 | phosphatase activity | 3.96E-04 | 13 |
|  | GO:0016758 | transferase activity, transferring hexosyl groups | 4.07E-04 | 14 |
|  | GO:0017171 | serine hydrolase activity | 1.14E-03 | 10 |
|  | GO:0008238 | exopeptidase activity | 1.90E-03 | 9 |
|  | GO:0008194 | UDP-glycosyltransferase activity | 6.87E-03 | 8 |

### Orthologs groups and orphan genes

We then searched the conservation and diversification of proteins among different *Phyllosticta* species from genome-scale. The protein orthology analysis identified 7261 orthologous groups existed in all the three *Phyllosticta* species, constituting the core gene set of *Phyllosticta* (Fig. 2). To find if any protein might be under positive selection, the dN/dS ratios for predicted proteins in a pairwise comparison between *P. citriasiana* and the other two *Phyllosticta* species were calculated. However, all gene show signs of purifying selection (dN/dS<1). 2040 genes in *P. capitalensis* have no orthologs in the other two species (Fig. 2), suggesting that these genes might play roles in constructing the endophytic relationship of *P. capitalensis* with its host. *P. citriasiana* and *P. citricarpa* encoded 371 and 262 species-specific proteins, respectively (Fig. 2), suggesting that these genes might be related to the host-specific pathogenicity. To know the functions of the genes in those three gene sets, we annotated them using the eggNOG database. However, the results showed that the majority of genes in each group encoded proteins without well-characterized domains and very few sequences can be assigned to the GO terms (Table S9-12).

**Fig. 2.**
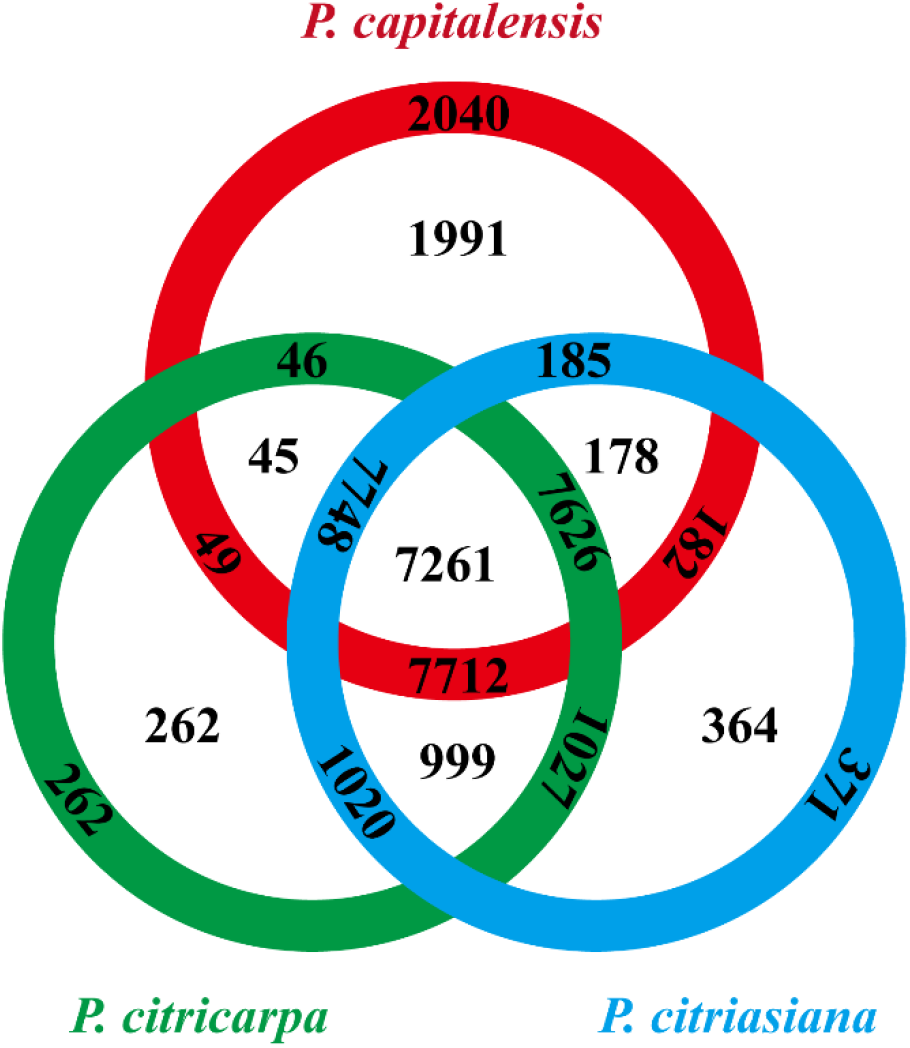
Numbers of orthologous groups that are unique to each isolate, specific to two isolates, and common to all three *Phyllosticta* isolates. Corresponding gene numbers are indicated in the outer ring.

We are then curious about if the distribution of CAZymes and secreted proteins might differ among different gene sets. We found that the *P. capitalensis* orphan genes contain only one CAZyme gene which encodes the AA3 family of cellobiose dehydrogenase while the other two species’ orphan genes contain no CAZyme gene (Table S13). However, the case for the secreted proteins is much different. *P. capitalensis*, *P. citricarpa*, and *P. citriasiana* contain 75, 8 and 17 species-specific secreted proteins, respectively (Table S13). These results indicate that *Phyllosticta* species have formed lineage-specific sets of orphan genes which might have a potential role in species diversification. Although functions of most orphans are unknown, the secreted proteins (potential effectors) are likely the essential factors of host specialization and they might be good candidates for future functional characterization in distinct *Phyllosticta* species.

### Secondary metabolite gene clusters

We identified 23 secondary metabolites (SM) biosynthesis clusters in the three *Phyllosticta* species (Table S14). These clusters are comprised of 3 NRPS clusters, 5 PKS clusters, 4 terpene clusters, 1 terpene-NRPS cluster, 1 PKS-NRPS cluster, and 9 clusters do not fit into any category (Table S14). Of them, cluster C9 contains all *Alternaria solani* genes involved in alternapyrone synthesis, suggesting that these *Phyllosticta* species have the potential to synthesize alternapyrone or its derivatives (Fujii et al., 2005). Most SM clusters (21) are well conserved among the three *Phyllosticta* species while 2 SM clusters of them showed whole cluster gain and loss polymorphisms or gene content polymorphisms. Cluster C7 was present in *P. citricarpa* and *P. citriasiana* but absent from *P. capitalensis*, indicating that this cluster might be lost in *P. capitalensis* or gained in the common ancestor of *P. citricarpa* and *P. citriasiana*. Meanwhile, Cluster C7 in *P. citricarpa* possesses another 3 genes while *P. citriasiana* contains a ~11 Kb region encoding no proteins, showing gene content polymorphisms (Fig. 3). SM cluster C23 showed two gene content polymorphisms. One is that the *P. capitalensis* has an additional 4 genes between orthologous gene OG0649 and OG0648. The other is *P. citriasiana* lost two genes, of which gene OG7537 encodes the backbone of this cluster (Fig. 3). A following tBLASTn analysis against the *P. citriasiana* genome confirmed the loss of these two genes. This gene content polymorphism was most likely generated through a genomic deletion event, rendering the SM gene cluster nonfunctional.

**Fig. 3.**
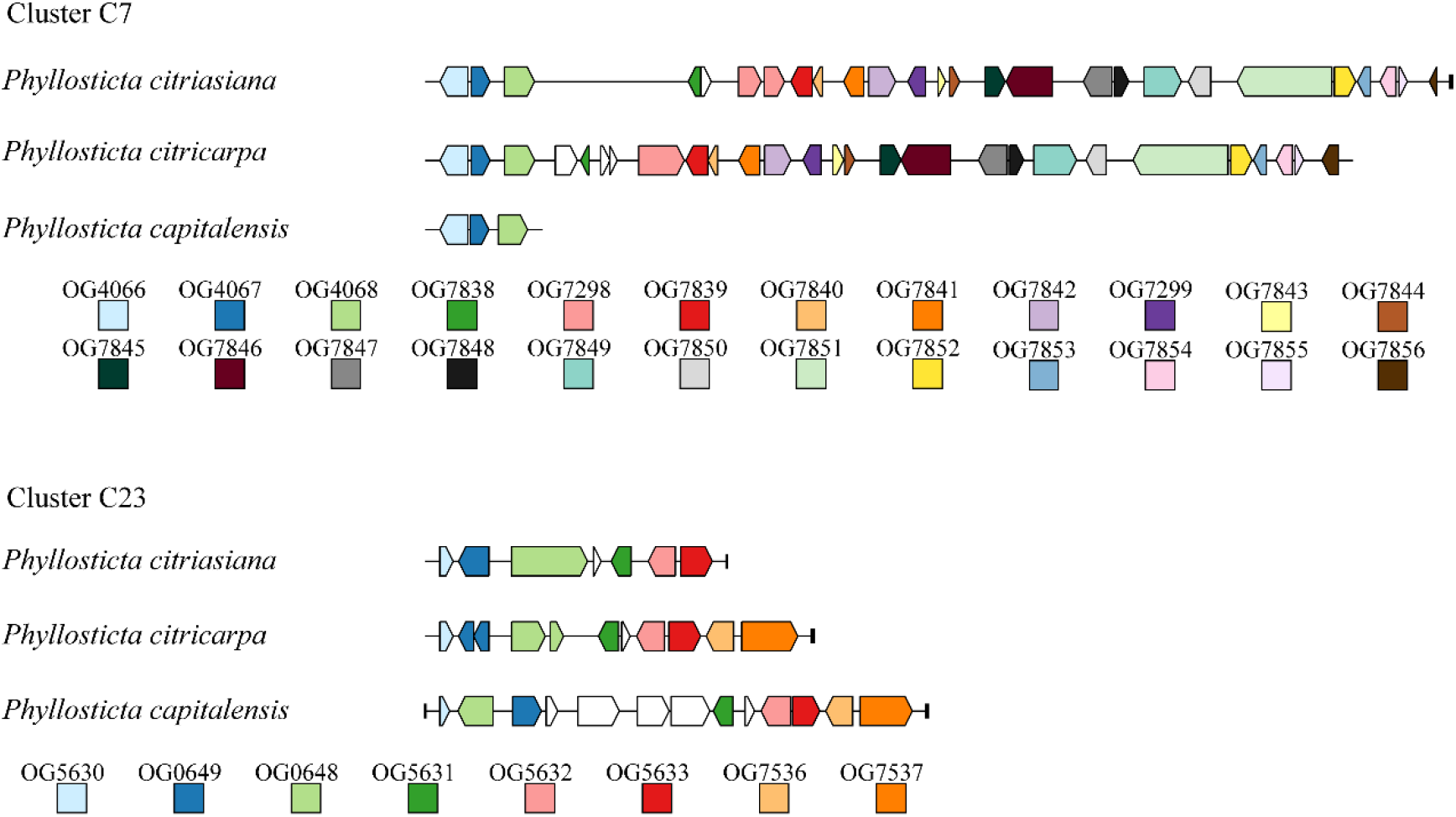
Structural variations of secondary metabolic (SM) gene cluster C7 and C23 among *Phyllosticta* species associated with citrus. For each SM cluster, orthologues among different species are marked with the same color. Genes marked by white lack orthologues in other species. The short black vertical line indicates the end of the contig.

Secondary metabolites, especial fungal toxins, are believed to be involved in the pathogenicity of many plant pathogenic fungal species and can be described as potential virulence factors. Previously, a handful of secondary metabolites from the citrus pathogen *P. citricarpa* were identified and characterized. Of them, a new dioxolanone, phenguignardic acid butyl ester, showed low phytotoxic activity in citrus leaves and fruits (at a dose of 100 μg) (Savi et al., 2019). However, the involvement of this compound in the formation of citrus black spot disease needs to be further addressed. In this study, we observed the major structural variation of two SM clusters among different *Phyllosticta* species, therefore, distinct corresponding metabolites are expected. However, if they are involved in the host specialization are not known. So, future investigations and elucidations of secondary metabolic mechanisms in *Phyllosticta* species and their functions involved in plant-fungal interactions will be of great significance.

## Conclusions

In this study, we sequenced the genome of *P. citriasiana*, the causal agent of the pomelo tan spot, generating a high-quality reference genome assembly and provide an overview of the genome structure of this important pathogen. We performed comparative genomics analysis to reveal overall high similarities in sequence identity and gene content among *P. citriasiana*, *P. citricarpa*, and *P. capitalensis*, reflecting the phylogenetic and ecological relatedness of these species associated with citrus. Our data also highlighted several striking differences in the constitution of secretomes, species-specific genes, and secondary metabolite gene clusters, which might contribute to the formation of fungal diversity. However, it is yet to be determined how a *Phyllosticta* species emerged as a pathogen of alternate hosts. These data would be valuable in the future investigation of the driving forces of fungal host switch, in population genomic studies for identification of haplotypes and alleles, and in identifying effectors that may function in infection of citrus plants.

## Author Contributions

MW and HL conceived the study. All authors analyzed the data. MW wrote the paper. All authors reviewed the manuscript.

